# Directed Evolution of a Plant Immune Receptor for Broad Spectrum Effector Recognition

**DOI:** 10.1101/2024.09.30.614878

**Authors:** Ellen Y. Rim, Oscar D. Garrett, Alexander J. Howard, Yejin Shim, Yuanyuan Li, Jonathan E. Van Dyke, Ryan C. Packer, Nguyen Ho, Rashmi Jain, Valley Stewart, Savithramma P. Dinesh-Kumar, James H. Notwell, Pamela C. Ronald

## Abstract

Rapid development of immune receptors that protect crops from emerging pathogens is a critical challenge ^1,2^. While novel immune receptors that recognize previously undetected pathogen effectors could provide protection against a wider range of pathogens, engineering such receptors has been constrained by the low throughput and speed of *in planta* testing. We established yeast surface display as a high throughput platform to recapitulate plant immune receptor-ligand interactions and evolve new binding capabilities. Using this directed evolution platform, we engineered the ligand binding domain of the rice immune receptor Pik-1 to recognize diverse effectors from the fast-evolving fungal pathogen *Magnaporthe oryzae.* Our approach yielded Pik-1 ligand binding domains with affinity for variants of the *M. oryzae* effector Avr-Pik that previously escaped detection by known rice alleles of Pik-1, with *in planta* assays confirming functional recognition of these effectors. Additional rounds of mutagenesis and selection led to a Pik-1 domain that binds all tested Avr-Pik variants as well as the evolutionarily divergent effector AvrPiz-t. These results demonstrate the potential of directed evolution to engineer immune receptors with new-to-nature recognition of a wide range of pathogen-derived ligands and accelerate development of broad spectrum resistance in crops.

## Main Text

Pathogens and host organisms are locked in an evolutionary competition: the host immune system strives to detect attacks while pathogens evolve to escape detection. At the molecular level, host immune recognition often hinges on direct interaction between a pathogen-derived molecule and an immune protein. Pathogens can accumulate mutations that escape such interactions, with their short lifecycles and large populations giving them an advantage. Directed evolution harnesses high throughput screening of synthetic protein sequences and effectively compresses the evolutionary timescale. Therefore, directed evolution can shift the balance in favor of the host by accelerating the development of immune proteins that recognize a wide array of pathogen-derived molecules, including those that evade detection by naturally evolved immune proteins.

Directed evolution can benefit plants in a unique manner. Unlike animals with their adaptive immune systems, plants carry hundreds to thousands of genome-encoded immune receptors. Many of these receptors are activated by direct binding to effectors, pathogen-secreted molecules that facilitate infection ^1^. Developing novel immune receptors that can bind previously unrecognized effectors, therefore, can protect crops against a broader spectrum of pathogens. Increasing global temperatures, which are predicted to enhance pathogen spread and reduce plant immunity, add urgency to development of such receptors ^2,3^. While gain-of-function random mutagenesis screens have led to improved immune receptors, their utility has been constrained by the limited throughput of *in planta* testing ^4–7^. Our goal was to establish a rapid, high throughput method to explore vast synthetic sequence spaces and identify variants with broad-spectrum ligand recognition. We chose Pik-1, an intracellular NOD-like receptor (NLR) in rice with a well-characterized effector binding domain ^8–10^, as our engineering target.

Rice blast, a fungal disease caused by *Magnaporthe oryzae*, leads to annual loss of rice enough to feed 60 million people ^11,12^. Introduction of immune receptors that protect against *M. oryzae* into crops presents an effective strategy to combat this global threat. Pik-1 recognizes the *M. oryzae* effector Avr-Pik through its heavy metal associated (HMA) domain, which has been co-opted from rice endogenous targets of Avr-Pik ^8,13,14^. Due to its modular nature, the Pik-1 HMA domain has been engineered extensively to alter ligand binding properties while retaining Pik-1 function in immunity ^15–19^. While Pik-1 recognition of Avr-Pik effectively triggers disease resistance, some *M. oryzae* strains harbor Avr-Pik variants with point mutations that escape detection by all known Pik-1 alleles; some strains lack Avr-Pik altogether. Eighty-five percent of sequenced rice-infecting *M. oryzae* strains that lack Avr-Pik variants harbor AvrPiz-t, a sequence divergent effector that belongs to the same family as Avr-Pik (*Magnaporthe* AVRs and ToxB-like, or MAX effector family) ^20,21^.

Coupling heterologous single cell expression with rounds of directed evolution (Fig. 1a), we engineered Pik-1 HMA domains that bind all known Avr-Pik alleles. Pik-1 containing these engineered domains triggered an immune response when co-expressed with Avr-Pik effectors in plant assays. We then further optimized the HMA domain for affinity to AvrPiz-t, reasoning that an engineered immune receptor that binds both sets of effectors would lead to broad spectrum resistance. Our results reveal that directed evolution can yield plant immune receptors with the new-to-nature ability to recognize multiple sequence divergent ligands. These findings open a new avenue for enhancing crop resistance to emerging pathogens.

**Fig. 1.**
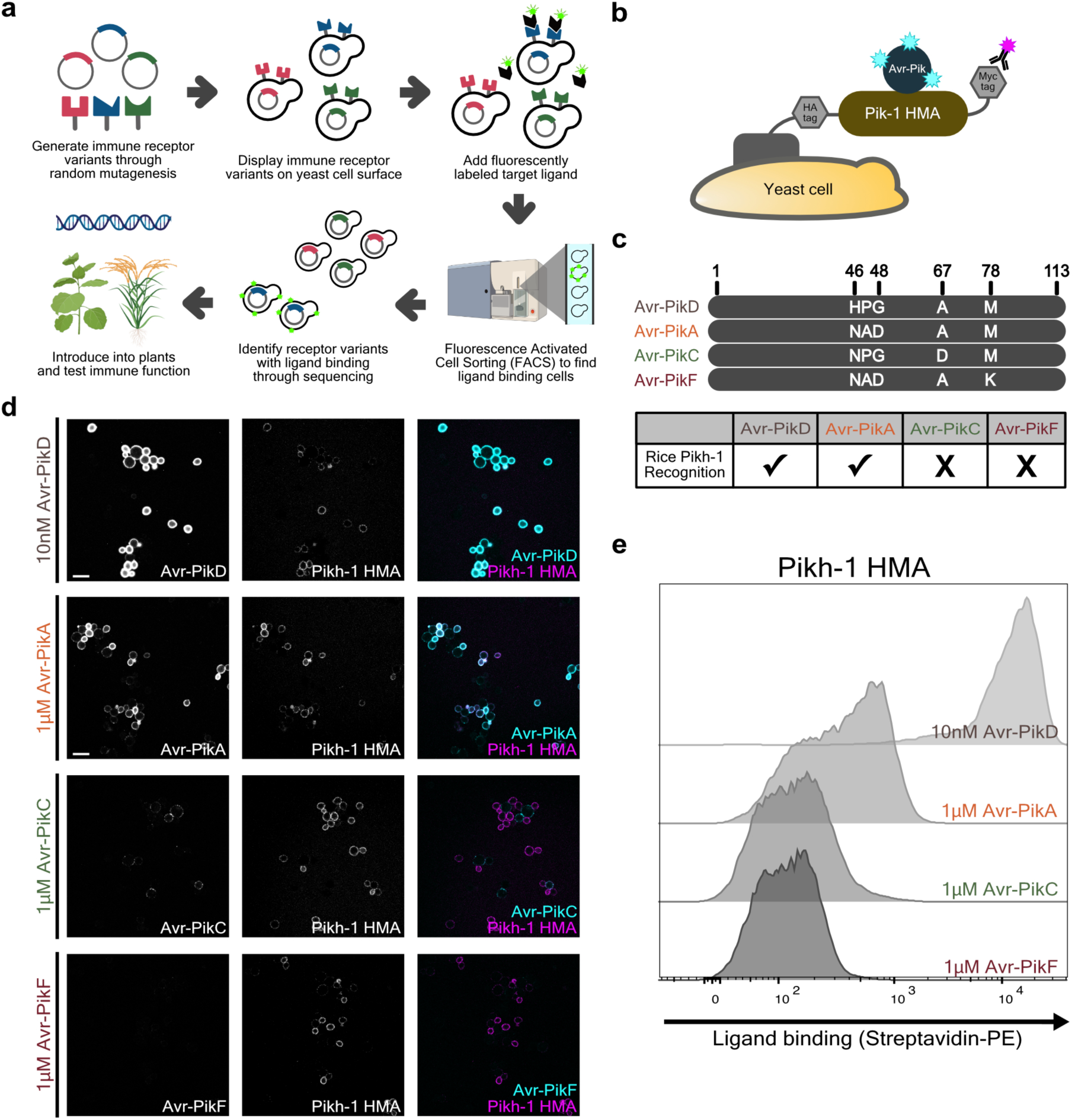
Yeast surface display of the rice receptor heavy metal associated (HMA) domain recapitulates its endogenous ligand binding properties. **a,** A schematic representation of directed evolution of a plant immune receptor with desired binding properties through yeast surface display. **b,** A schematic representation of Pikh-1 yeast surface display. Pikh-1 HMA domain expression and its interaction with target Avr-Pik are assessed through two-color fluorescence detection. **c,** Allelic variants of the *M. oryzae* effector Avr-Pik. Rice receptor Pikh-1 elicits immune response in the presence of Avr-PikD and Avr-PikA, but not Avr-PikC and Avr-PikF. **d,** Immunofluorescence images of yeast cells expressing Pikh-1 HMA domain. Left: biotin-labeled effector binding at specified concentrations detected through Alexa Fluor 488-conjugated streptavidin. Middle: HMA domain expression detected through anti-Myc tag binding to Alexa Fluor 568. Right: merged images of effector binding (cyan) and receptor expression (magenta). Scale bar, 10µm. **e,** Flow cytometry quantification of the interaction between yeast surface-displayed Pikh-1 HMA domain and biotin-labeled Avr-Pik variants.

### Plant immune receptor-ligand interactions are recapitulated in a yeast expression platform

We first set out to test whether the Pik-1 HMA domain could retain its effector interaction properties when expressed on *Saccharomyces cerevisiae*. In yeast surface display, recombinant proteins fused to a cell wall protein are displayed on the surface, allowing screening of up to ∼10^9^ variants in one experiment. This approach has identified proteins with desired ligand interactions, including novel receptor domains that bind pathogen-derived molecules or antibodies with antitumor or pathogen neutralizing activities ^22–24^. We expressed three well-characterized rice genome-encoded alleles of the HMA domain of Pik-1 (Pikp-1, Pikm-1, and Pikh-1) tethered to the yeast Aga2p cell wall protein and flanked by epitope tags (Fig. 1b).

We tested whether effector binding specificity and relative affinity were recapitulated by yeast surface displayed Pik-1 HMA domains (Fig. 1c). Fluorescence imaging of Myc epitope-stained yeast cells showed expression of Pik-1 HMA domains on the cell surface and their interaction with biotin-labeled effectors (Fig. 1d and Extended Data Fig. 1a). The Pikh-1 allele exhibits one of the widest effector recognition ranges. The HMA domain of Pikh-1 showed strong affinity for Avr-PikD, moderate affinity for Avr-PikA, but no significant interaction with Avr-PikC and Avr-PikF variants of the effector, reproducing the effector binding properties of this allele in rice ^25^. Flow cytometry analysis of Pik-1 HMA domain-expressing yeast cells recapitulated these results (Fig. 1e and Extended Data Fig. 1b,c). These results validated that our yeast expression platform reflects the specificity and affinity of Pik-1 immune receptor-ligand interactions.

### Directed evolution of Pik-1 HMA domain variants that bind previously unrecognized ligands

We then sought to engineer yeast displayed Pikh-1 HMA domain to bind previously unrecognized effectors through directed evolution. We first subjected the HMA domain to random mutagenesis through error prone PCR. Mutagenized amplicons were introduced into yeast cells to generate a library of Pikh-1 HMA domain variants. Colony dilution estimated the library to contain 2.2×10^7^ variants and sequencing a subset of the library showed that the average number of amino acid mutations was 2.1 over the 78aa in the effector binding domain, with the majority of variants containing one to three amino acid changes (Extended Data Fig. 2a and Supplementary Table 1).

To identify Pikh-1 HMA domain variants that bind the previously unrecognized effectors Avr-PikC and Avr-PikF, we performed four rounds of FACS selection (Fig. 2a). Starting with the library, we exposed yeast cells displaying Pikh-1 HMA domain variants to either biotin-labeled Avr-PikC or Avr-PikF and selected those showing the highest affinity for the target effector and robust HMA domain expression. In subsequent selection rounds, variants were incubated with lower concentrations of the target effector alternating between Avr-PikC or Avr-PikF, with the goal of identifying variants that bind both effectors with high affinity (Fig. 2a). In each selection round we sorted through enough cells to oversample the population diversity by five- to ten-fold to avoid losing unique variants, selecting cells in the top 0.5-1% for target binding and HMA domain expression to carry onto the next round (Extended Data Fig. 2b).

**Fig. 2.**
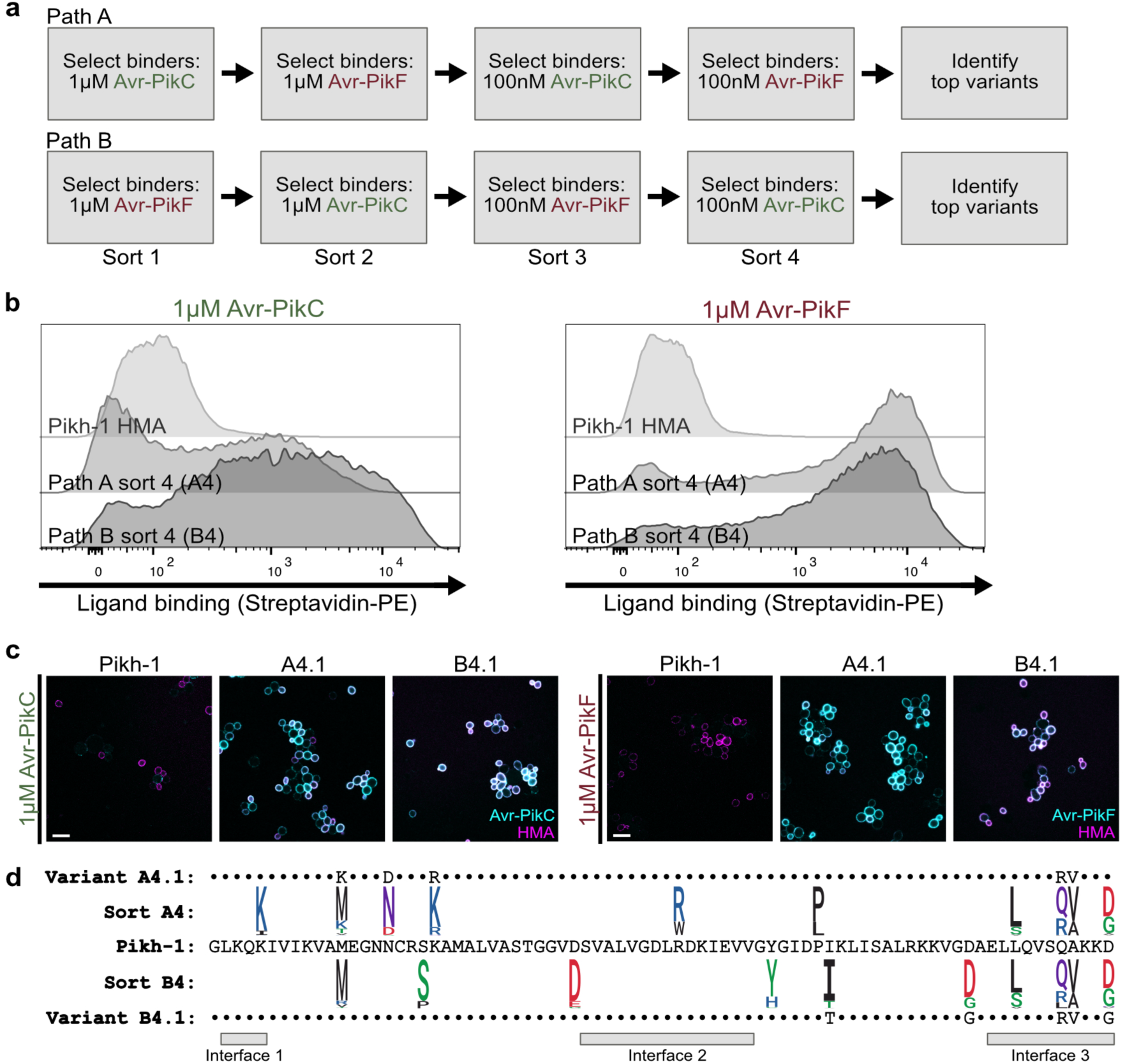
Rounds of FACS selection identify Pik-1 HMA domain variants with affinity for novel ligands. **a**, Directed evolution workflow to develop binding to both Avr-PikC and Avr-PikF. **b**, Flow cytometry quantification of the interaction between Pikh-1 HMA domains and 1µM Avr-PikC or 1µM Avr-PikF. Compared to the wildtype Pikh-1 HMA domain (top), variants collected at the end of selection path A (middle) and path B (bottom) show increased affinity for Avr-PikC (left) and Avr-PikF (right). **c**, Immunofluorescence images of yeast cells expressing Pikh-1 HMA domains in the presence of 1µM Avr-PikC (left) or 1µM Avr-PikF (right). Merged images show biotin-labeled effector binding detected through Alexa Fluor 488-conjugated streptavidin (cyan) and receptor expression detected through anti-Myc tag binding to Alexa Fluor 568 (magenta) of the wildtype Pikh-1, A4.1, and B4.1 HMA domains. Scale bar, 10µm. **d**, Amino acid sequences of the wildtype Pikh-1 HMA domain and variants A4.1 and B4.1 enriched at the end of selection paths A and B, respectively. Sequence logos depict the 10 most frequently mutated residues after sort 4 of each selection path (red: acidic, blue: basic, black: hydrophobic, purple: neutral, green: polar). Interfaces known to interact with Avr-Pik are indicated below ^10,16^.

Following four rounds of selection, variants gained binding to both Avr-PikC and Avr-PikF at the population level regardless of the order in which they were exposed to the two effectors (Fig. 2b). Illumina sequencing of these populations showed that several variants had increased in representation over selection rounds (Supplementary Table 2). Of these, variants we named A4.1 and B4.1 showed the greatest enrichment relative to the starting library at the end of selection path A and selection path B, respectively (Supplementary Table 3).

These variants indeed exhibited high affinity for both Avr-PikC or Avr-PikF (Fig. 2c). Flow cytometry and imaging analysis indicated that their affinities for Avr-PikC and Avr-PikF were comparable to or exceeded the binding affinity required for Pik-1-induced *in planta* immune response (Extended Data Figs. 1b and 2c). Importantly, A4.1 and B4.1 retained the ability to bind all effectors recognized by the wildtype Pikh-1 HMA domain: Avr-PikA, Avr-PikD, and Avr-PikE (Extended Data Fig. 2d). A4.1 and B4.1 also showed binding to a novel allele of Avr-Pik we identified in *M. oryzae* isolates TW-6-2-2-B-1 and JS-10-6-1-2 and named Avr-PikJ (Extended Data Fig. 2d). An Avr-Pik-like effector with ∼60% sequence identity to Avr-Pik, Avr-PikL2A, was previously identified in rice-infecting *M. oryzae* (Extended Data Fig. 2e) ^26^. Variants A4.1 and B4.1 bound Avr-PikL2A whereas the wildtype Pikh-1 HMA domain failed to do so (Extended Data Fig. 2d).

A4.1 and B4.1 harbor five substitutions, with two of them (Q74R and A75V of the HMA domain sequence) shared by both variants (Fig. 2d). A75V is also present in interface 3 of the natural allele Pikm-1 and related variants, which is thought to contribute extensively to the interaction with Avr-Pik effectors in these alleles (Extended Data Fig. 3a) ^27^. However, the rest of the mutations that conferred broad spectrum Avr-Pik binding in A4.1 and B4.1 had not been observed in rice genome-encoded or previously engineered alleles of Pik-1. Therefore, these two Pikh-1 HMA domain variants gained the ability to bind all known alleles of Avr-Pik through unique, new-to-nature mutations.

### Directed evolution of Pik-1 HMA domain variants with broad spectrum ligand binding

Directed evolution can yield a receptor that binds allelic variants of its native ligand, but can additional rounds of selection lead to affinity for a more divergent ligand? To test this, we targeted AvrPiz-t. While Avr-Pik and AvrPiz-t both belong to the MAX family of *M. oryzae* effectors characterized by their β-sandwich structures, they exhibit no detectable sequence similarity (Fig. 3a and Extended Data Fig. 2e). Avr-Pik and AvrPiz-t are recognized by two distinct rice genome-encoded immune receptors and interact with different sets of host proteins to suppress immunity, highlighting different evolutionary trajectories and functions of the two effectors ^13,20,28–30^. AvrPiz-t is present in 85% of rice-infecting *M. oryzae* that lack all alleles of Avr-Pik; 193 out of 201 rice-infecting isolates with genomes available on NCBI harbor either Avr-Pik or AvrPiz-t. Its divergence from Avr-Pik, high conservation in *M. oryzae* strains that lack Avr-Pik, and role in facilitating pathogen infection rendered AvrPiz-t both a good test case and a practical target for broad spectrum *M. oryzae* recognition by an engineered receptor.

**Fig. 3.**
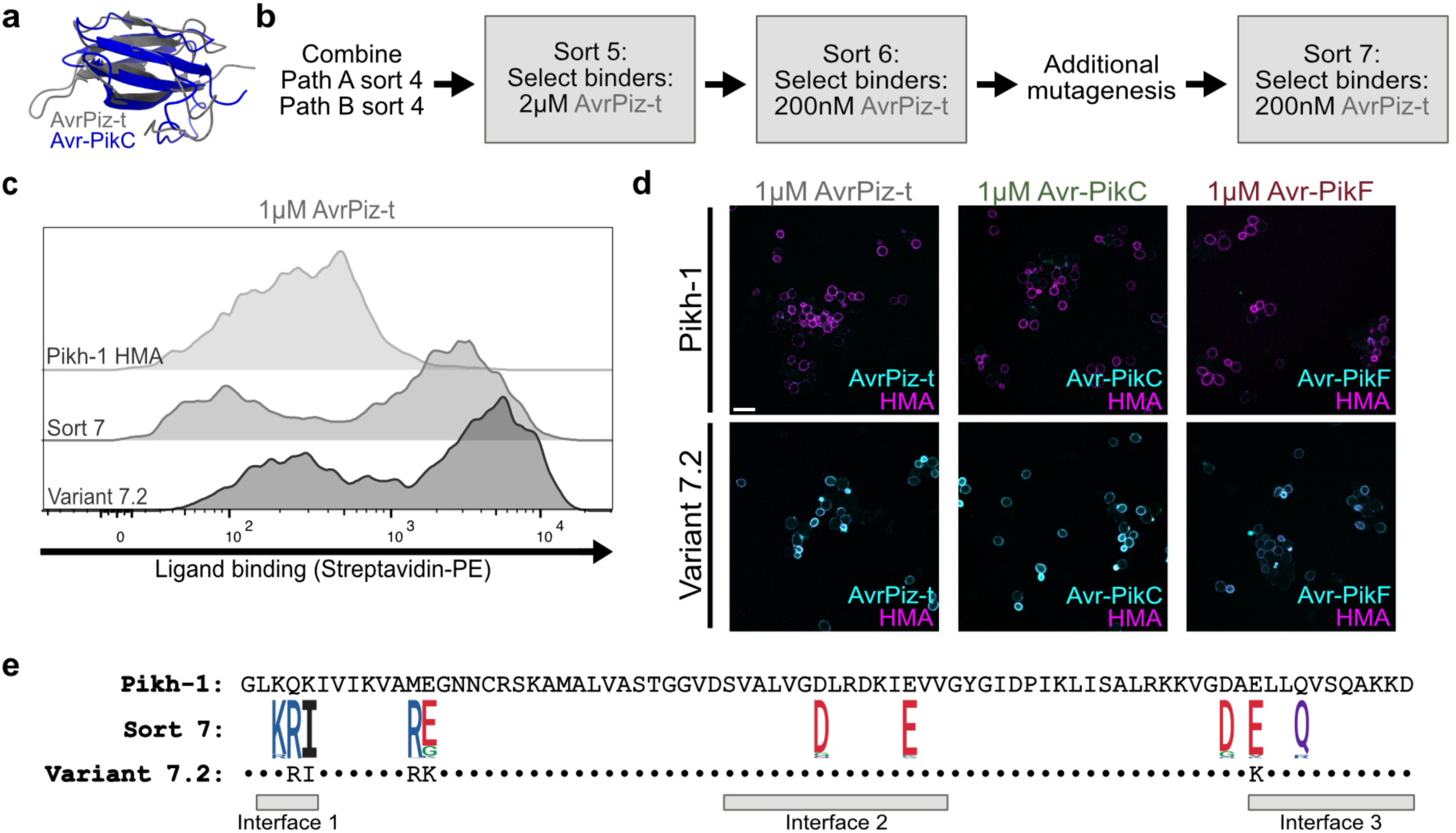
Additional FACS selection identifies Pik-1 HMA domain variants with affinity for an evolutionarily divergent ligand. **a**, Overlay of crystal structures of Avr-PikC (blue, PDB: 7A8X) and AvrPiz-t (grey, PDB: 2LW6). **b**, Directed evolution workflow to develop binding to AvrPiz-t. Variants were selected for binding to AvrPiz-t at decreasing concentrations, with an additional round of random mutagenesis to enhance library diversity. **c**, Flow cytometry quantification of the interaction between Pikh-1 HMA domains and 1µM AvrPiz-t. Compared to the wildtype Pikh-1 HMA domain (top), variants collected at the end of sort 7 (middle) and selected variant 7.2 (bottom) show increased affinity for AvrPiz-t. **d**, Immunofluorescence images of yeast cells expressing Pikh-1 HMA domains in the presence of 1µM AvrPiz-t (left), 1µM Avr-PikC (middle), or 1µM Avr-PikF (right). Merged images show biotin-labeled effector binding detected through Alexa Fluor 488-conjugated streptavidin (cyan) and receptor expression detected through anti-Myc tag binding to Alexa Fluor 568 (magenta) of the Pikh-1 wildtype (top) and variant 7.2 (bottom) HMA domains. Scale bar, 10µm. **e**, Amino acid sequences of the wildtype Pikh-1 HMA domain and variant 7.2 are shown. Sequence logo depicts the 10 most frequently mutated residues after sort 7 (red: acidic, blue: basic, black: hydrophobic, purple: neutral, green: polar). Interfaces known to interact with Avr-Pik are indicated below ^10,16^.

To identify Pikh-1 HMA domain variants that bind AvrPiz-t in addition to Avr-PikC and Avr-PikF, we pooled variants collected after four rounds of selection through paths A and B. These variants were subjected to two subsequent rounds of selection to identify those with affinity for decreasing concentrations of AvrPiz-t (Fig. 3b). As top HMA variants identified at the end of the selection process did not exhibit strong binding to AvrPiz-t, we further diversified the post-sort 6 pool through error-prone PCR. This new library was estimated to contain 2.1×10^7^ variants through colony dilution, with an average of 4.4 mutations over the 78aa in the effector binding domain (Extended Data Fig. 3b). Variants collected after one additional round of selection with 200nM AvrPiz-t showed enhanced affinity for AvrPiz-t at the population level (Fig. 3c and Extended Data Fig. 3c). We evaluated Pikh-1 HMA variants that were significantly enriched following sort 7 relative to their frequencies in the post-sort 6 mutagenized library (Supplementary Table 3). Among these, variant 7.2 demonstrated strong binding to AvrPiz-t as well as Avr-PikC and Avr-PikF in confocal imaging and flow cytometry analyses (Fig. 3c,d and Extended Data Fig. 4a). Additionally, 7.2 showed affinity for all tested alleles of Avr-Pik in confocal imaging (Extended Data Fig. 4b).

The HMA domain variant with three (Q4R, K5I, and M12R) of the substitutions in 7.2 was present at low frequencies after four rounds of selection for Avr-PikC and Avr-PikF binding (Supplementary Table 2). This then became the predominant variant following selection rounds 5 and 6 for AvrPiz-t binding. However, these changes alone were insufficient to confer robust affinity for Avr-PikC, Avr-PikF, and AvrPiz-t (data not shown). The additional E13K and E68K substitutions in variant 7.2 appear to account for improved binding to all three ligands. Therefore, iterative cycles of stringent selection and diversity enrichment through random mutagenesis yielded a HMA domain variant capable of binding all Avr-Pik alleles and AvrPiz-t.

### Recognition of target ligands by engineered Pik-1 immune receptors in *N. benthamiana*

Binding interactions between the rice Pik-1 and *M. oryzae* Avr-Pik can be recapitulated in the model plant *Nicotiana benthamiana* ^10^. Interaction between *Agrobacterium tumefaciens-* delivered Pik-1 and Avr-Pik triggers immune activity and cell death at the site of *A. tumefaciens* infiltration (Fig. 4a) ^10^. We transiently expressed full-length Pikh-1 harboring wildtype or engineered HMA domain, its helper receptor Pikh-2 required for immune function, and various Avr-Pik effectors in *N. benthamiana*. Despite plant-to-plant variability in cell death severity, especially near the ligand recognition threshold, A4.1 or B4.1 consistently triggered cell death in response to Avr-PikC and Avr-PikF (Fig. 4b,c). The 7.2 HMA variant induced strong cell death with Avr-PikF and showed small but significant (p < 0.005, two-tailed Mann-Whitney test) improvement over wildtype HMA with AvrPiz-t (Fig. 4d,e). However, 7.2 did not enhance Avr-PikC recognition despite showing binding in yeast-based assays (Fig. 4d,e).

**Fig. 4.**
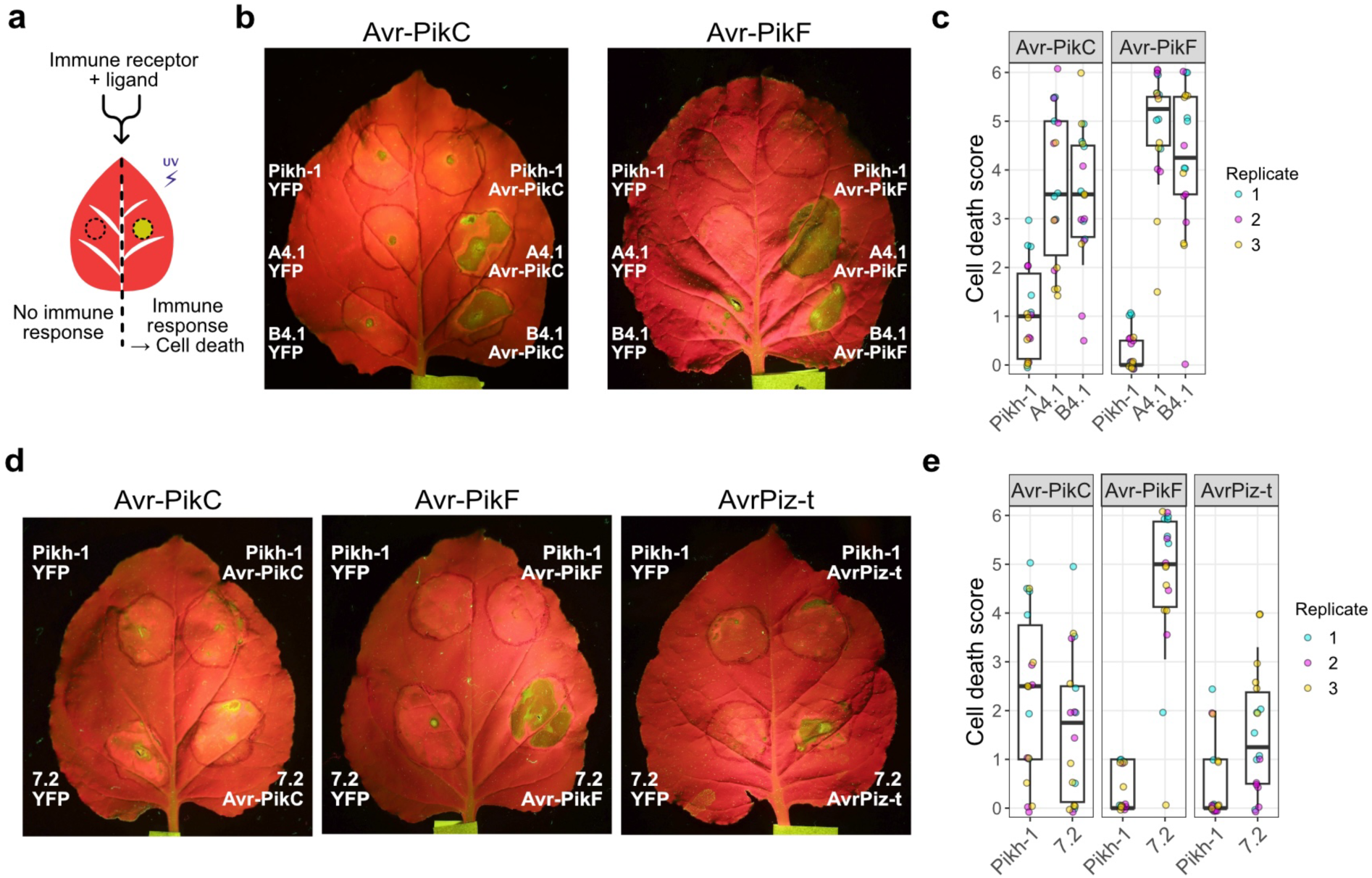
Recognition of target ligands by engineered Pik-1 immune receptors in *N. benthamiana*. **a**, A schematic representation of cell death assay in *N. benthamiana* to assess immune receptor function. **b**, Representative UV images of cell death assay. Pikh-1 harboring wildtype, A4.1, or B4.1 HMA domain was co-expressed with Avr-PikC (left leaf) or Avr-PikF (right leaf). YFP was co-expressed as a negative control. **c**, Quantification of cell death induced by Pikh-1 harboring wildtype, A4.1, or B4.1 HMA domain scored on a 0-6 scale. Center line denotes median value; box edges delineate 25^th^ and 75^th^ percentiles; whiskers extend from 10^th^ to 90^th^ percentiles. Each receptor-effector pair was tested in three independent experiments with six plants per experiment. **d**, Representative UV images of cell death assay. Pikh-1 harboring wildtype or 7.2 HMA domain was co-expressed with Avr-PikC (left leaf), Avr-PikF (middle leaf), or AvrPiz-t (right leaf). YFP was co-expressed as a negative control. **e**, Quantification of cell death induced by Pikh-1 harboring wildtype or 7.2 scored on a 0-6 scale. Center line denotes median value; box edges delineate 25^th^ and 75^th^ percentiles; whiskers extend from 10^th^ to 90^th^ percentiles. Each receptor-effector pair was tested in three independent experiments with six plants per experiment.

Changes to NLR effector binding domains can sometimes lead to autoactivity and trigger immune activation without the cognate effector ^15^. The engineered variants did not induce cell death in the absence of a target effector, demonstrating their specific activity (Fig. 4b,d, Extended Data Fig. 5b,c). Therefore, Pikh-1 variants evolved to bind previously unrecognized ligands triggered immune activation to target ligands in *N. benthamiana*. We note that transferring robust recognition of a more divergent set of ligands proved more challenging.

## Discussion

Our results demonstrate the first successful application of high throughput directed evolution to engineer a plant immune receptor domain that can bind previously unrecognized ligands. Lab-evolved Pikh-1 HMA domain variants displayed binding to all tested *M. orzyae* effectors in the Avr-Pik family as well as a sequence divergent effector, AvrPiz-t. An immune receptor that binds evolutionarily divergent effectors with distinct roles in pathogen infection is predicted to confer broad spectrum resistance in crops. If functionally validated in rice for immune function, such a receptor will likely maintain its effectiveness over time barring mutation or loss of expression of all target effectors in the pathogen.

Multiple evolutionary paths led to expanded effector perception in our experiments. A4.1 and B4.1 were obtained from different selection schemes and shared two out of their five amino acid substitutions. Yet both variants showed affinity for all known alleles of the effector Avr-Pik. Variant 7.2 additionally gained the ability to recognize AvrPiz-t through a yet different set of five synergistic changes. Substitutions in these engineered variants revealed residues relevant for effector recognition outside of the interfaces identified in receptor-effector crystal structures^10,15,25,27^.

When tested in *N. benthamiana*, engineered variants A4.1 and B4.1 activated immune responses to Avr-PikC and Avr-PikF, effectors that evade detection by known natural alleles of Pik-1. For variant 7.2, which was further engineered to bind AvrPiz-t, improvements in immune activation did not fully correlate with the binding strength observed in yeast assays for some effectors. 7.2 likely binds AvrPiz-t through a different interface than Avr-Pik, as demonstrated for HMA domains that bind sequence divergent MAX effectors ^31–33^. The substitutions in 7.2 that enable AvrPiz-t binding may affect conformational changes required for Pikh-1 activation, particularly near ligand recognition thresholds. These results highlight both the challenge and importance of validating engineered protein functions in plant systems.

Plant immune receptor engineering has largely relied on grafting ligand binding domains or subsets thereof from genome-encoded variants and testing chimeric receptors in individual plants. While effective, this approach is labor intensive and requires extensive structural knowledge of the receptor-effector interface, which is not available yet for most plant immune receptors ^34^. Directed evolution in a heterologous system can expand effector recognition spectra of immune receptors at a faster pace even without prior structural knowledge. For instance, a loss-of-function random mutagenesis screen performed in yeast can quickly identify the receptor domain required for interaction with a cognate effector. A subsequent gain-of-function screen, as described here, searches for receptor variants with affinity for previously unrecognized effectors.

Enhancing effector recognition through substitutions identified through high throughput screening offers additional advantages over introducing exogenous immune receptors or replacing large receptor segments. These small changes are more amenable to current crop genome editing technologies. Introducing precise amino acid changes into a genome-encoded receptor such as Pik-1 through prime editing, for example, presents a feasible path to develop and deploy improved crops (31, 32). Our results demonstrate how high throughput directed evolution can explore large synthetic sequence spaces to develop immune recognition of divergent effectors, a function that seemed initially far-fetched. We have established directed evolution as a strategy to develop broad spectrum resistance and protect crops from fast evolving pathogens.

---

**Extended Data Fig. 1.**
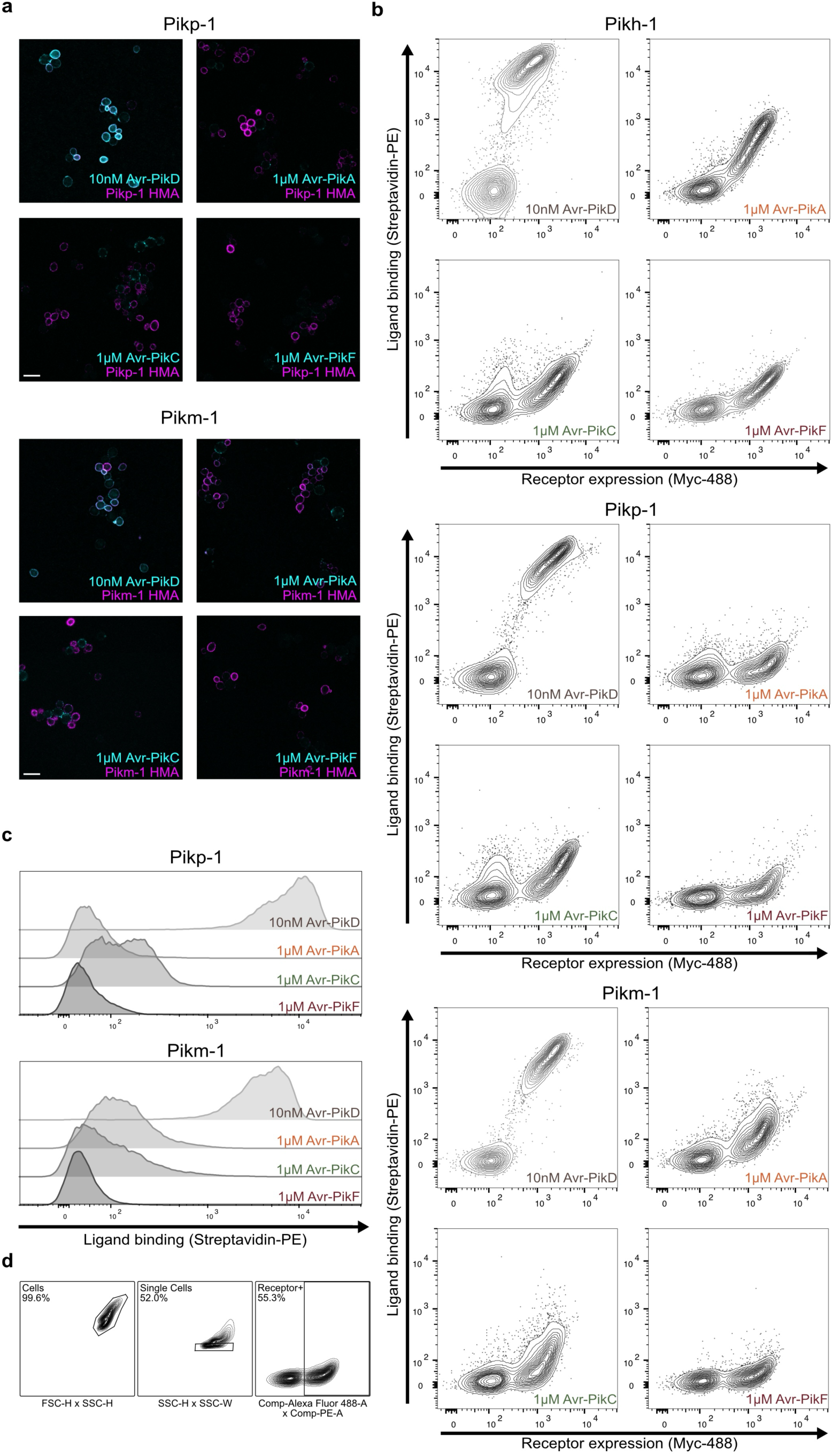
Yeast surface-displayed Pik-1 alleles recapitulate their endogenous ligand binding properties. **a**, Immunofluorescence images of yeast cells expressing Pikp-1 or Pikm-1 HMA domain in the presence of Avr-Pik variants. Merged images show biotin-labeled effector binding detected through Alexa Fluor 488-conjugated streptavidin (cyan) and HMA domain expression detected through anti-Myc tag binding to Alexa Fluor 568 (magenta). Scale bar, 10μm. **b**, Flow cytometry quantification of Pikh-1, Pikp-1, and Pikm-1 HMA domain surface expression and interaction with Avr-Pik variants shown in bivariate contour plots. Clusters of cells that do not express the target protein ^35^ do not exhibit Avr-Pik binding. **c**, Flow cytometry quantification of Pikp-1 and Pikm-1 HMA domain interaction with Avr-Pik variants shown in univariate histograms. **d**, Exemplary flow cytometry ancestry plots showing the gating strategy applied to histograms in this study, as described in the Methods section. Plots from Pikm-1 HMA domain interaction with 1μM Avr-PikF are shown as an example.

**Extended Data Fig. 2.**
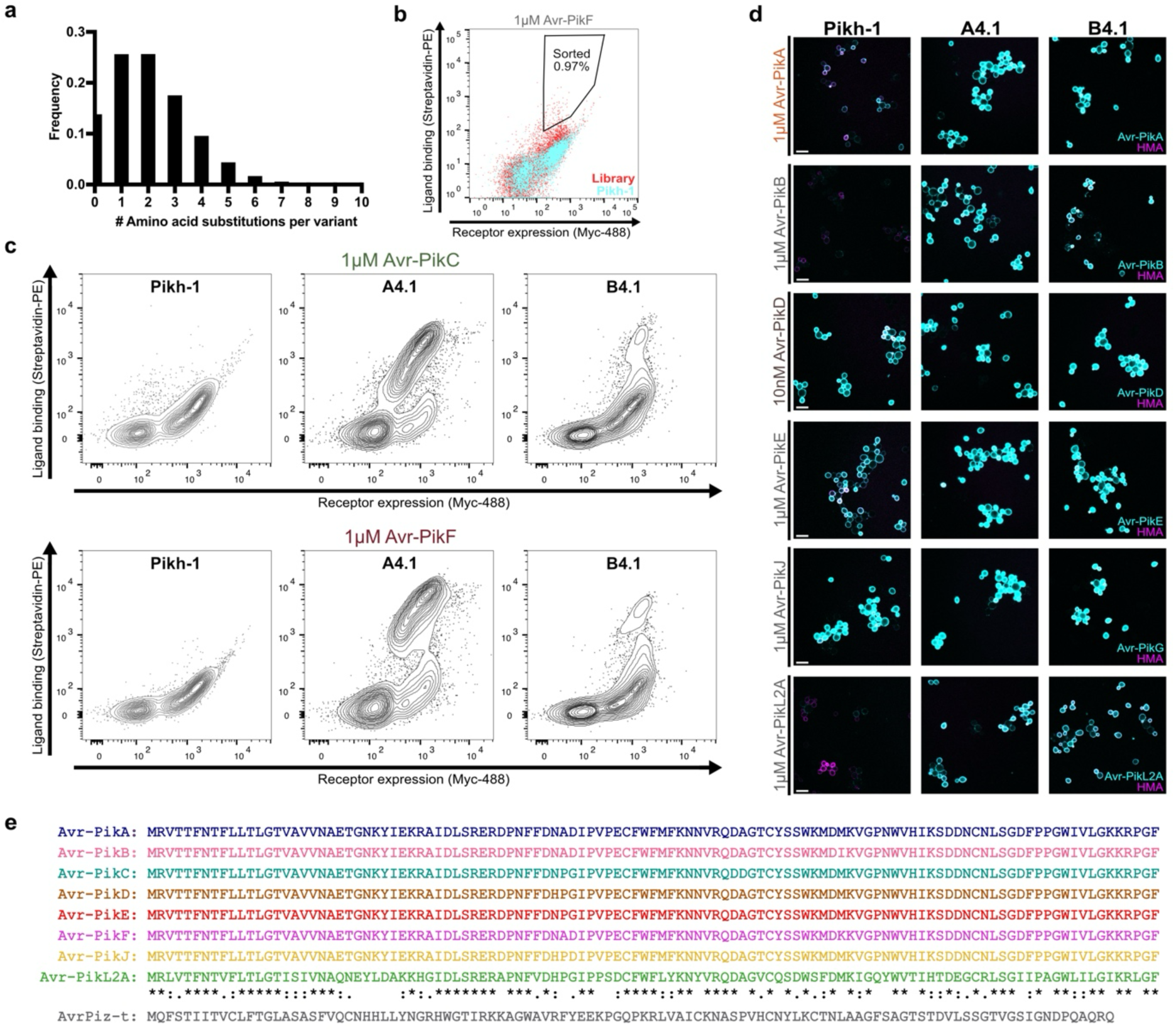
Iterative selection leads to Pik-1 HMA domain variants with improved affinity for Avr-PikC and Avr-PikF. **a**, Frequency of variants harboring the specified number of substitutions over the 78-amino acid HMA domain in the library of 2.2×10^7^ Pikh-1 HMA domain variants. **b**, Bivariate dot plot of Pikh-1 HMA domain surface expression and interaction with 1μM Avr-PikF showing an exemplary FACS selection window. Cyan dots indicate cells expressing the wildtype Pikh-1 HMA domain, and red dots indicate cells expressing Pikh-1 HMA domain variants in the library. **c**, Flow cytometry quantification of Pikh-1, A4.1, and B4.1 HMA domain surface expression and interaction with 1μM Avr-PikC or Avr-PikF shown in bivariate contour plots. **d,** Immunofluorescence images of yeast cells expressing Pikh-1, A4.1, or B4.1 HMA domain in the presence of each effector at specified concentrations. Merged images show effector binding (cyan) and expression (magenta) of HMA domains. Scale bar, 10μm. **e**, Amino acid sequences of Avr-Pik and AvrPiz-t effectors from *Magnaporthe oryzae*.

**Extended Data Fig. 3.**
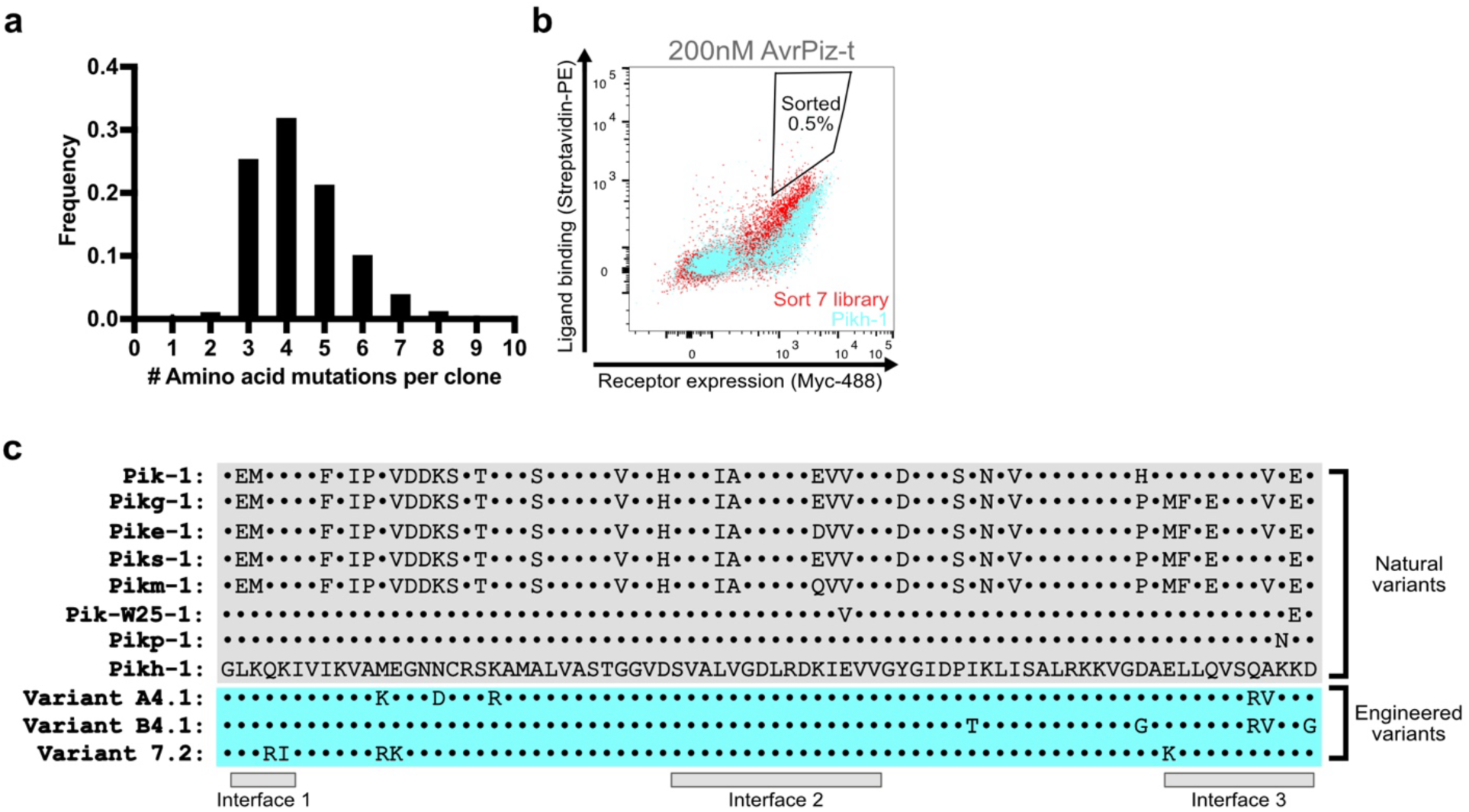
Characterization of the second library and sequences of natural and engineered Pik-1 variants. **a**, Frequency of variants harboring the specified number of substitutions over the 78-amino acid HMA domain in the library of 2.1×10^7^ Pikh-1 HMA domain variants following the second round of mutagenesis. **b,** Bivariate dot plot of Pikh-1 HMA domain surface expression and interaction with 200nM AvrPiz-t showing an exemplary FACS selection window. Cyan dots indicate cells expressing the wildtype Pikh-1 HMA domain and red dots indicate cells expressing Pikh-1 HMA domain variants in the library following second round of mutagenesis. **c**, Amino acid sequences of Pik-1 HMA domains from rice alleles Pik-1, Pikg-1, Pike-1, Piks-1, Pikm-1, Pik-W25-1, Pikp-1, and Pikh-1 and engineered variants A4.1, B4.1, and 7.2. Interfaces known to interact with Avr-Pik are indicated below ^10,16,36,37^.

**Extended Data Fig. 4.**
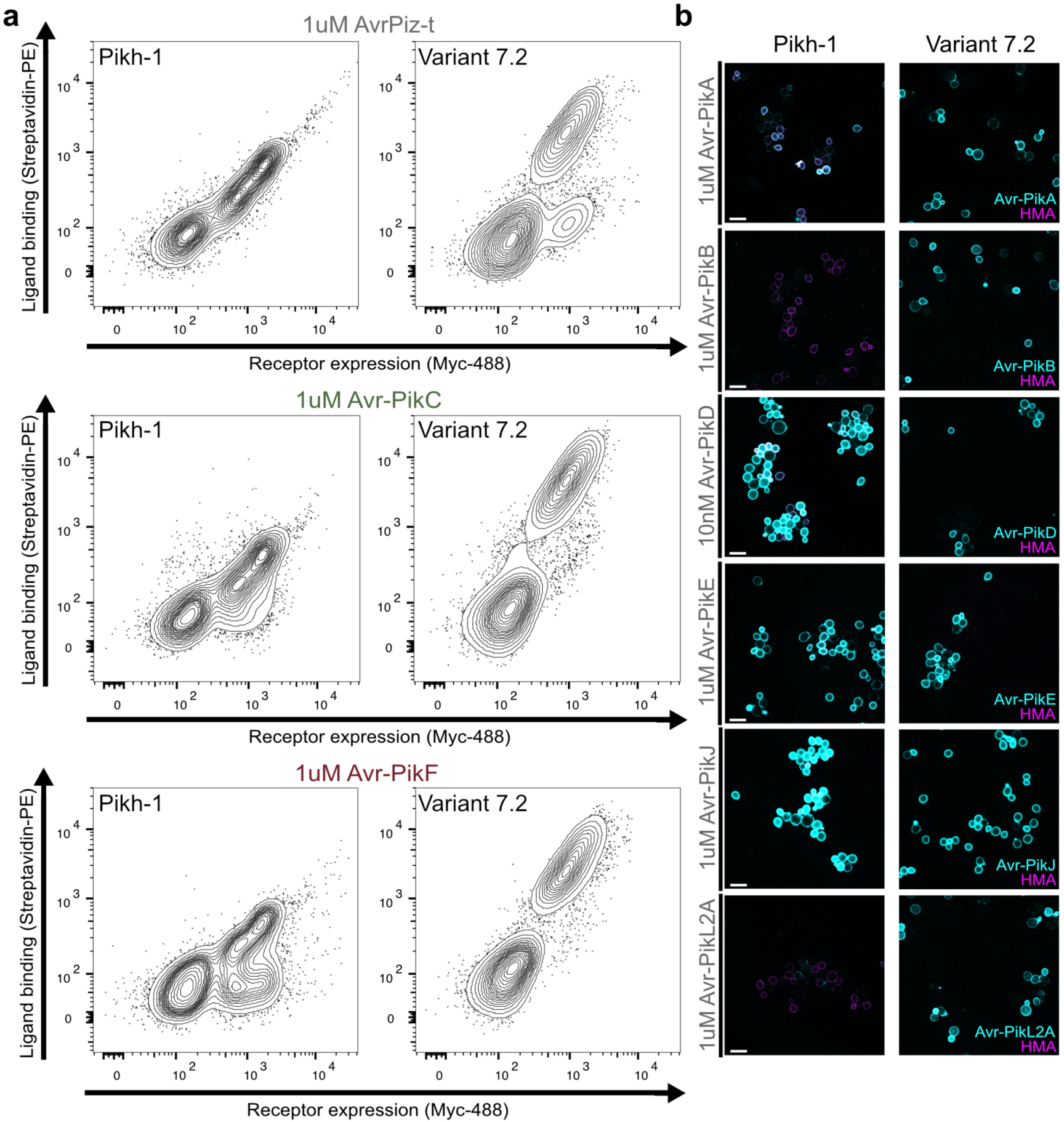
Additional rounds of selection lead to a Pik-1 HMA domain variant with affinity for Avr-Pik and AvrPiz-t. **a**, Flow cytometry quantification of Pikh-1 and 7.2 HMA domain surface expression and interaction with 1μM AvrPiz-t, Avr-PikC, or Avr-PikF shown in bivariate contour plots. **b**, Immunofluorescence images of yeast cells expressing Pikh-1 or 7.2 HMA domains in the presence of each effector at specified concentrations. Merged images show effector binding (cyan) and expression (magenta) of HMA domains. Scale bar, 10μm.

**Extended Data Fig. 5.**
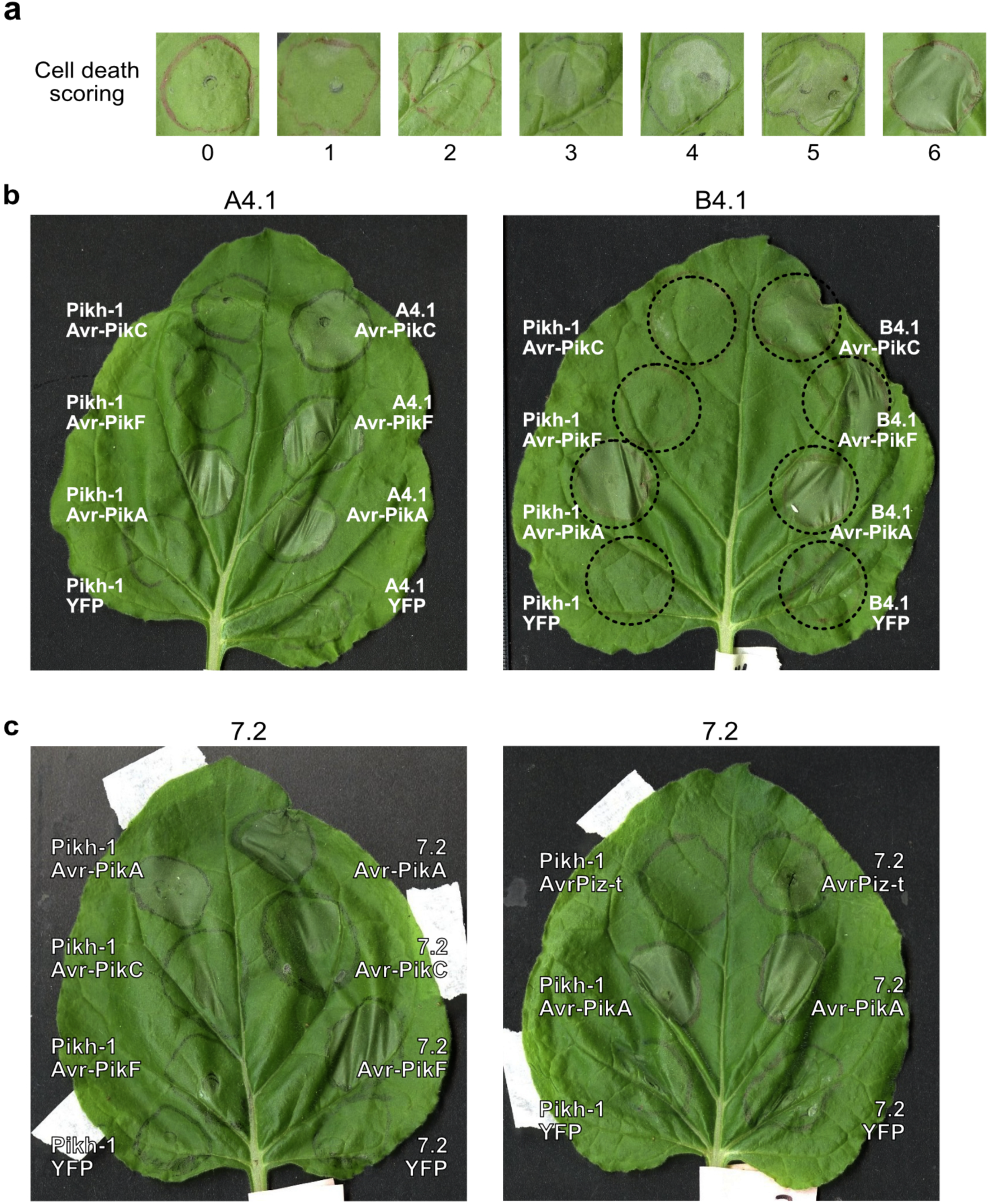
Engineered Pik-1 immune receptors induce cell death response in *N. benthamiana*. **a**, Representative images of *N. benthamiana* cell death scores on the 0-6 scale used in this study. **b**, Representative images of cell death assay. Pikh-1 harboring wildtype, A4.1, or B4.1 HMA domain was co-expressed with Avr-PikC, Avr-PikF, Avr-PikA (positive control), or YFP (negative control). **c**, Representative images of cell death assay. Pikh-1 harboring wildtype or 7.2 HMA domain was co-expressed with Avr-PikC, Avr-PikF, AvrPiz-t, Avr-PikA (positive control), or YFP (negative control).

## Methods

### DNA constructs

For yeast surface display: Yeast codon optimized sequences of rice Pik-1 HMA domains (aa186-263) was synthesized by Twist Biosciences and cloned into the yeast display vector pCTcon2 using NheI/BamHI restriction sites. pCTcon2 (Addgene plasmid #41843) was a gift from Dane Wittrup ^38^. For E. coli expression of effectors: pOPIN-GG vectors encoding Avr-PikD/A/C tagged with 6xHis and GB1 were gifts from Adam Bentham and Mark Banfield ^39^. Vectors encoding Avr-PikE and Avr-PikF were generated by site-directed mutagenesis of Avr-PikD and Avr-PikA, respectively. Avr-PikB and Avr-PikJ sequences were synthesized by Twist Biosciences and cloned into pOPIN-GG using NotI/BamHI restriction sites. AvrPiz-t and AvrPikL2A sequences PCR amplified from pUC57-Kan_GY11_00131271 (Addgene plasmid #121355) and pUC57-Kan_FR13_APikL2 (Addgene plasmid #123648), which were gifts from Sophien Kamoun ^40^, and Gibson cloned into pOPIN-GG. For cell death assay in *N. benthamiana*: Rice Pikh2 sequence was synthesized by Twist Biosciences and cloned into the binary vector pGFPGUSPlus (Addgene plasmid #64401), which was a gift from Claudia Vickers ^41^, using SacI/XbaI restriction sites. CaMV 35S promoter followed by rice Pikh1 sequence was synthesized by Twist Biosciences and introduced into pGFPGUSPlus Pikh2 using SbfI/BstEII restriction sites. This resulted in a binary vector with Pikh2 and Pikh1 with wildtype or engineered HMA domain, each flanked by CaMV 35S promoter and Nos terminator. Avr-PikA, Avr-PikC, Avr-PikF, AvrPiz-t, and P19 were introduced into pEarleyGate104 ^42^ using PspXI/PacI restriction sites. This resulted in a vector with the coding sequence flanked by CaMV 35S promoter and OCS terminator. Sequences of all primers and DNA constructs are listed in Supplementary Table 4.

### Library preparation

Pikh-1 HMA domain in the yeast surface display vector pCTcon2 was subjected to error prone PCR following published library generation protocols ^38,43,44^. Briefly, Pikh-1 HMA domain sequence was amplified using Taq polymerase in mutagenic reactions with 1µM to 2µM of 8- oxo-dGTP (TriLink N-2034) and dPTP (TriLink N-2037) for 15 cycles. These reactions were pooled and further amplified to yield ∼20µg DNA. pCTcon2 vector DNA was digested with SalI, NheI, and BamHI at 37°C overnight. 4µg of digested vector was mixed with 12µg of the mutagenized insert and precipitated using Pellet Paint following the manufacturer’s protocol (Millipore 69049). Mutagenized library of yeast surface-displayed Pikh-1 HMA domain was introduced into yeast strain EBY100 (ATCC MYA-4941) via electroporation ^24,44,45^. 100 mL of YPD media [1% (w/v) yeast extract, 2% peptone, 2% D-glucose] was inoculated with stationary phase EBY100 to an OD_600_ of 0.3. Cells were incubated at 30°C with shaking at 225rpm until OD_600_ of 1.6 was reached. Cells were pelleted at 3000 xg for 3 minutes, washed twice with 50mL cold MilliQ water, once with 50mL cold electroporation buffer [1M sorbitol, 1mM CaCl_2_], and incubated in 20mL of lithium buffer [0.1M LiAc, 10mM DTT] for 30 minutes at 30°C with shaking at 225rpm. Cells were resuspended in 1mL electroporation buffer. 400µL of the electrocompetent EBY100 was mixed with 4µg linearized vector and 12µg mutagenized insert. Following electroporation using a Bio-Rad GenePulser Xcell at 2.5 kV and 25 µF in a 2mm cuvette, cells were suspended in 10mL of 1:1 YPD:1M sorbitol and incubated at 30°C for 1 hour. Cells were centrifuged and resuspended in 100mL SDCAA [2% (w/v) D-glucose, 0.54% disodium phosphate, 0.86% monosodium phosphate, 0.5% casamino acids, 0.67% yeast nitrogen base]. Serial dilution from 1/100 to 1/100,000 estimated library size of 2.2 × 10^7^.

### Yeast surface display

Single clones of wildtype or mutated Pikh-1 HMA domain were introduced into yeast strain EBY100 using Frozen-EZ Yeast Transformation II kit (Zymo Research) following the manufacturer’s protocol. Transformed yeast was grown on SDCAA agar plates [SDCAA with 1.5% agar and 18.2% sorbitol] at 30°C for 2-3 days. Individual colonies were picked and grown in 5mL SDCAA at 30°C with shaking at 225rpm for 16 hours. To induce expression of Pikh-1 HMA domain on yeast, SDCAA cultures were centrifuged at 1800 xg for 3 minutes and washed with SGCAA [2% (w/v) galactose, 0.54% disodium phosphate, 0.86% monosodium phosphate, 0.5% casamino acids, 0.67% yeast nitrogen base]. Cells were resuspended in SGCAA to a final concentration of 1 × 10^7^ cells/mL and grown at 25°C with shaking at 225 rpm for 24 hours.

### Confocal microscopy

Approximately 10^6^ yeast cells were pelleted at 3000 xg at 4°C for 3 minutes and resuspended in 100µL PBSA [1% bovine serum albumin in PBS pH 7.4]. After adding a biotinylated effector to the specified concentration, cells were incubated rotating at 25°C for 1.5 hours. In the last 30 minutes, 1:100 Myc tag rabbit monoclonal antibody (Cell Signaling 2278) was added to detect Pikh-1 HMA domain expression. Cells were pelleted and washed with 500µL PBSA and resuspended in 100µL PBSA with 1:100 anti-rabbit IgG secondary antibody Alexa Fluor 568 conjugate (Thermo Fisher A-11010) and 1:100 streptavidin Alexa Fluor 488 conjugate (Thermo Fisher S11223). Secondary staining was performed on ice in the dark for 30 minutes. After a final wash with 500µL PBSA, cell pellet was resuspended in 30µL PBSA. 6-7µL of the stained cell suspension was mounted on a glass slide and covered with 22mm x 22mm #1 thickness coverslip. Samples were imaged on Leica TCS SP8 confocal microscope with 63x oil immersion lens and excitation with 488nm and 552nm lasers.

### Fluorescence-activated cell sorting

Published yeast library FACS protocols were modified for selection of Pikh-1 HMA domain variants that bind target effectors ^43,44,46,47^. For the initial sort, 10^8^ yeast cells were pelleted at 3000 xg at 4°C for 3 minutes and resuspended in 1mL PBSA and divided into five 1.7mL tubes. After adding a biotinylated effector to the specified concentration, cells were incubated rotating at 25°C for 1.5 hours. In the last 30 minutes, 1:100 Myc-tag rabbit monoclonal antibody (Cell Signaling 2278) was added to detect Pikh-1 HMA domain expression. Cells were pelleted and washed with 1mL PBSA and resuspended in 200µL PBSA with 1:100 anti-rabbit IgG secondary antibody Alexa Fluor 488 conjugate (Thermo Fisher A-11008) and 1:100 streptavidin PE conjugate (Thermo Fisher S866). Secondary staining was performed on ice in the dark for 30 minutes. After a final wash with 1mL PBSA, cell pellet was resuspended in 4mL PBSA and passed through 35µm strainer mesh into FACS tubes (Corning 352235). Cells expressing the wildtype Pikh-1 HMA domain and exposed to 10nM Avr-PikD were used to prepare unstained, 488 or PE single stained, and double stained control samples. Top 1% cells with the high PE and 488 fluorescence values, representing target effector binding and Pikh-1 HMA domain expression, respectively, were sorted into SDCAA. Approximately 10^6^ sorted cells were grown in 50mL SDCAA at 30°C with shaking at 225 rpm until saturation. These cells were frozen down in 15% glycerol in SDCAA or passaged in SDCAA for use in subsequent rounds of FACS. Each subsequent round started with at least tenfold the number of cells collected in the previous round to ensure adequate sampling of the variant diversity. Reagent to detect target effector binding alternated between streptavidin PE conjugate and anti-biotin PE conjugate (Thermo Fisher 50-169-246) to prevent selection for secondary reagent binding. Experiments were performed on Beckman Coulter Astrios EQ cell sorter and Becton Dickinson FACS Aria II cell sorter at the UC Davis Flow Cytometry Shared Resource. FACS data were analyzed and plots were generated using FlowJo software v10.9.0 (Becton Dickinson). To generate histograms, yeast cells were gated first using forward versus side scatter, then single cells were gated using side scatter height versus width. Receptor expressing cells gated using Alexa Fluor 488 area versus PE area were then plotted in a histogram, with same gates applied to all samples (Extended Data Fig. 1d).

### Yeast DNA extraction and sequencing

For yeast DNA extraction, Zymoprep Yeast Plasmid DNA Miniprep II kit (Zymo Research) was used following the manufacturer’s protocol. Briefly, >10^6^ yeast cells were grown in 2mL SDCAA at 30°C with shaking at 225rpm for 4 hours. Cells were pelleted at 16000 xg for 3 minutes and resuspended in 200µL digestion buffer with 5µL Zymolase. Following a 2-hour incubation at 37°C, cell lysate was prepared using lysis and neutralization buffers from Zymoprep Yeast Plasmid DNA Miniprep II kit. DNA was extracted using GenElute Plasmid Miniprep Kit (Sigma) and columns and eluted in 20µL MilliQ water. 50ng of the eluted DNA was used as template in a 100µL PCR reaction with 500nM pCTcon2 Illumina Fwd/Rev primers, 200µM dNTP, 3% DMSO, and 2 units of Phusion High Fidelity DNA polymerase (New England Biolabs) in 1x Phusion HF buffer. Reaction was initially heated to 98°C for 5 minutes then cycled 30 times at 98°C for 10 seconds, 60°C for 30 seconds, and 72°C for 1 minute, and finally extended at 72°C for 10 minutes, yielding 410bp amplicons containing the Pikh-1 HMA domain insert. PCR product was purified using DNA Clean & Concentrator Kit (Zymo Research) and eluted in 20µL MilliQ water. Up to 1µg of purified DNA was sent for library preparation, sequencing reaction, and initial bioinformatics analysis at AZENTA, Inc. DNA library was prepared with NEBNext Ultra DNA Library Prep Kit for Illumina (New England Biolabs) following the manufacturer’s protocol and sequenced using a 2x 250 paired-end (PE) configuration on an Illumina instrument. Raw sequence reads were trimmed of their adapters and nucleotides with poor quality using Trimmomatic v. 0.36. This returned 338000 reads per sample on average. After further filtering reads with indels or nonsense mutations, reads that encode unique Pikh-1 HMA domain amino acid sequences were counted. Pikh-1 HMA domain variants detected and their read counts in the starting library and FACS selected samples can be found in Supplementary Table 1 and Supplementary Table 2.

### Effector purification and biotinylation

Avr-Pik and AvrPiz-t effectors were expressed using pOPIN-GG vectors generated and shared by Bentham and colleagues ^39^. Effectors tagged with 6xHis and GB1 in pOPIN-GG were introduced into Shuffle T7 Express Competent E. coli (New England Biolabs C3029). Single colonies were grown in 5mL Terrific Broth [1.2% (w/v) tryptone, 2.4% (w/v) yeast extract, 0.94% (w/v) dipotassium phosphate, 0.22% (w/v) monopotassium phosphate, 0.4% (v/v) glycerol, 1mM magnesium sulfate] with 100µg/mL carbenicillin at 30°C with shaking at 150rpm for 20 hours. These cultures were used to inoculate 500mL fresh TB with 100µg/mL carbenicillin and grown at 30°C with shaking at 100rpm to an OD_600_ of 0.6-0.8. To induce protein expression, IPTG was added to 1mM for 18 hours at 20°C with shaking at 100rpm. Cells were harvested by centrifugation at 4000 xg at 4°C for 20 minutes and frozen at -80°C. Thawed cell pellet was mixed with 10mL lysis buffer [50mM HEPES pH 8.0, 500mM NaCl, 50mM glycine, 5% (v/v) glycerol, 30mM imidazole, 1 tablet EDTA-free protease inhibitor cocktail (Millipore Sigma 11873580001)], incubated on ice for 30 minutes, and sonicated with approximately 600J energy. Lysate was collected following centrifugation at 10000 xg at 4°C for 20 minutes. Effectors were affinity purified using Ni-NTA Spin Columns (Thermo Fisher 88224) following the manufacturer’s protocol and eluted in 500mM imidazole in lysis buffer. Samples were desalted using Spin Desalting Columns (Thermo Fisher 89892) and buffer exchanged into A4 buffer [10mM HEPES pH 7.4, 150mM NaCl] using Amicon Ultra-0.5 Centrifugal Filter Unit 10KDa (Millipore UFC501024) following manufacturers’ protocols. 6xHis and GB1 tag was cleaved by adding 20U/mg protein 3C Protease (Thermo Fisher 88946) and rotating for 16 hours at 4°C. Cleaved effectors were collected in the flow-through after binding to Ni-NTA Spin Columns, desalted, and buffer exchanged into PBS pH 8.0. Sample purity was confirmed in a Coomassie stained gel and BCA Protein Assay (Thermo Fisher 23225) measured effector amount of 200µg-1mg per prep. Effectors were biotin labeled by adding 40-fold molar excess NHS-PEG_4_-biotin (Thermo Fisher 21455) and rotating for 1 hour at 24°C. To quench the reaction, Tris pH 8.0 was added to 50mM final concentration and rotated for 30 minutes at 24°C. Excess reagents were eliminated from samples using G-10 Macro SpinColumns (Harvard Apparatus 74-3904). Biotin labeled and purified effectors were stored in -80**°**C until use in FACS or imaging experiments.

### Nicotiana benthamiana cell death assay

*N. benthamiana* plants were grown in a growth chamber with 16-hour light cycle with 150µmol/m^2^ light intensity and 24°C day and 22°C night temperatures. Plants were germinated and grown in Sunshine Mix #1 (Sun Gro Horticulture) and watered with 2:1:2 N:P:K macronutrient ratio fertilizer. *Agrobacterium tumefaciens* GV3101 carrying desired constructs were resuspended in infiltration media [10 mM MES pH 5.7, 10 mM MgCl_2_, 500 µM Acetosyringone]. Sequences of all DNA constructs are listed in Supplementary Table 4. *A. tumefaciens* carrying Pikh-2 and wildtype or engineered Pikh-1 (final OD600 0.6) was mixed with Avr-Pik, AvrPiz-t, or YFP (final OD600 0.6) and P19 (final OD600 0.2). The fourth leaf from the top of each five-week-old *N. benthamiana* was infiltrated using a needleless syringe and the plants were kept in the dark post infiltration ^48^. Leaves were collected at 3 days post infiltration for white light and UV autofluorescence imaging. Cell death response of each infiltrated spot was blindly scored using a 0-6 scale (Extended Data Fig. 5a) and two independent scores were averaged. Each engineered Pikh-1 was tested in three experiments with six plants per experiment.

### Data availability

All data generated or analyzed during this study are included in this article and its supplementary table files.

## Acknowledgments

We thank A. Bentham and M. Banfield for sharing pOPIN-GG vectors and for advice on protein expression and purification. This project was supported by the University of California Davis Flow Cytometry Shared Resource Laboratory with technical assistance from Bridget McLaughlin, Jonathan Van Dyke and Ashley Karajeh.

## Funding

Life Sciences Research Foundation – Simons Foundation (EYR)

National Institutes of Health MIRA 1R35GM148173 (PCR)

The Chan Zuckerberg Initiative (PCR)

National Science Foundation IOS-2139987 (SPD-K)

National Institutes of Health grant P30 CA093373 (Flow Cytometry Shared Resource)

National Institutes of Health NCRR C06-RR1208 (Flow Cytometry Shared Resource)

National Institutes of Health S10 OD018223 (Flow Cytometry Shared Resource)

National Institutes of Health S10 RR 026825 (Flow Cytometry Shared Resource)

James B. Pendleton Charitable Trust (Flow Cytometry Shared Resource)

Partially supported by the Joint BioEnergy Institute, U.S. Department of Energy, Office of Science, Biological and Environmental Research Program under Award Number DE-AC02-05CH11231 (PCR)

## Author contributions

Conceptualization: EYR, JHN, PCR

Formal Analysis: EYR, AJH, JHN

Funding acquisition: EYR, PCR

Investigation: EYR, ODG, AJH, YS, YL, RCP

Methodology: EYR, VJS, JEV

Project administration: EYR, PCR

Software: AJH, JHN, NH, RJ

Resources: YL, SPD

Supervision: PCR, JHN, SPD

Validation: EYR, ODG, YS

Visualization: EYR, AJH

Writing – original draft: EYR

Writing – review & editing: EYR, JHN, PCR

## Competing interests

E.Y.R. and P.C.R. are inventors on a provisional patent application (No. 63/692,024) related to this work.

## Notes

### Summary of Updates

The manuscript has been revised to include additional data - functional validation in planta.

