## Supplementary Data for "Directed Evolution of a Plant Immune Receptor for Broad Spectrum Effector Recognition"

**The PDF file includes:**

Extended Data Figs. 1 to 5

**Other Supplementary Information for this manuscript include the following:**

Supplementary Tables 1 to 4

**Supplementary Table 1.**
Variant sequences and read counts in the initial library.

**Supplementary Table 2.**
Variant sequences and read counts following sort 4 and sort 7.

**Supplementary Table 3.**
Log transformed enrichment scores following sort 4 and sort 7.

**Supplementary Table 4.**
Sequences of primers and constructs used in the study.

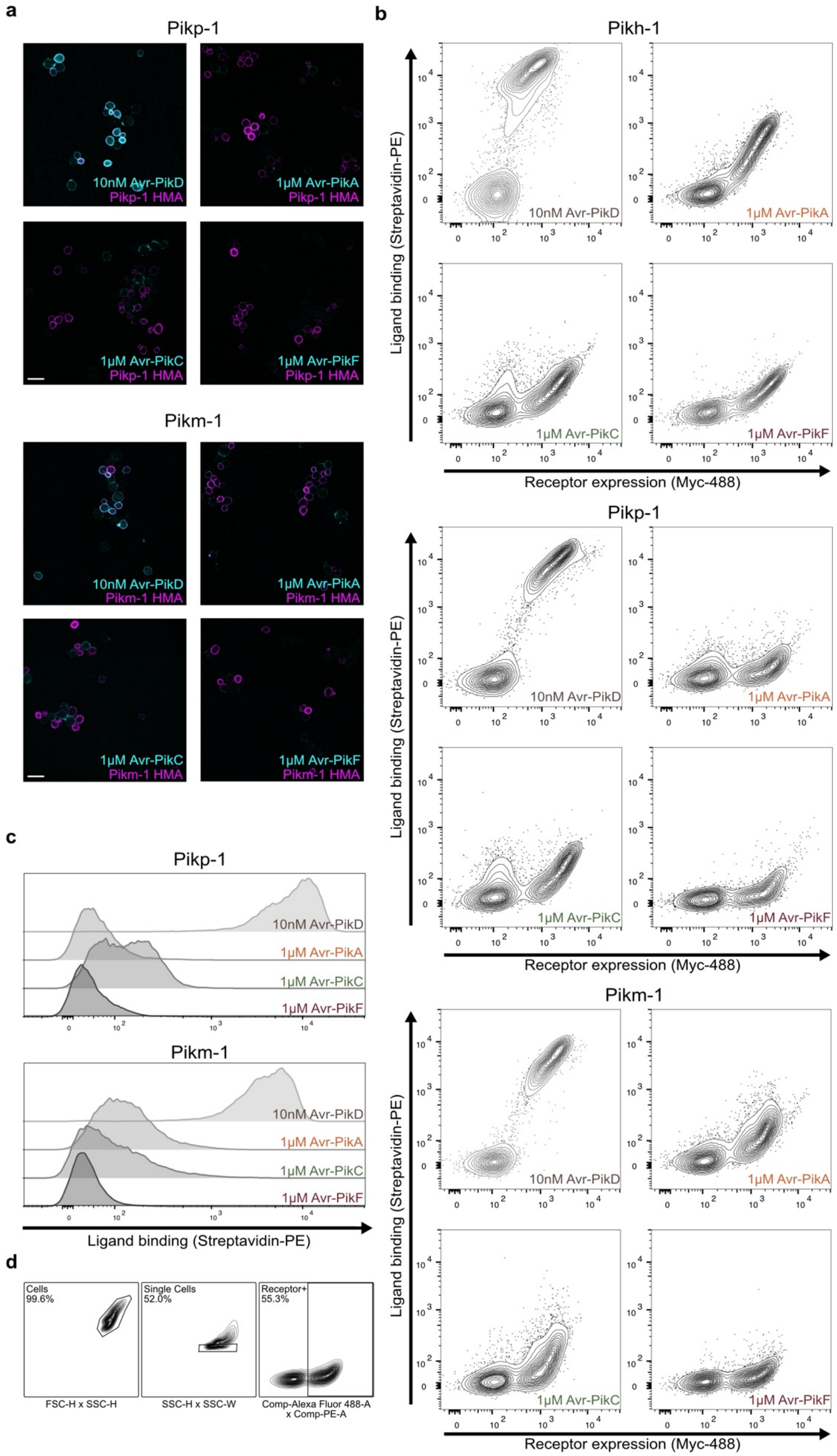
a
Pikp-1
10nM Avr-PikD
Pikp-1 HMA
1μM Avr-PikA
Pikp-1 HMA
1μM Avr-PikC
Pikp-1 HMA
1μM Avr-PikF
Pikp-1 HMA
Pikm-1
10nM Avr-PikD
Pikm-1 HMA
1μM Avr-PikA
Pikm-1 HMA
1μM Avr-PikC
Pikm-1 HMA
1μM Avr-PikF
Pikm-1 HMA
b
Pikh-1
Ligand binding (Streptavidin-PE)
Receptor expression (Myc-488)
10nM Avr-PikD
1μM Avr-PikA
1μM Avr-PikC
1μM Avr-PikF
Pikp-1
Pikm-1
c
Pikp-1
Pikm-1
10nM Avr-PikD
1μM Avr-PikA
1μM Avr-PikC
1μM Avr-PikF
Ligand binding (Streptavidin-PE)
d
Cells
99.6%
Single Cells
52.0%
Receptor+
55.3%
FSC-H x SSC-H
SSC-H x SSC-W
Comp-Alexa Fluor 488-A
x Comp-PE-A

**Extended Data Fig. 1. Yeast surface-displayed Pik-1 alleles recapitulate their endogenous ligand binding properties. a**, Immunofluorescence images of yeast cells expressing Pikp-1 or Pikm-1 HMA domain in the presence of Avr-Pik variants. Merged images show biotin-labeled effector binding detected through Alexa Fluor 488-conjugated streptavidin (cyan) and HMA domain expression detected through anti-Myc tag binding to Alexa Fluor 568 (magenta). Scale bar, 10μm. **b**, Flow cytometry quantification of Pikh-1, Pikp-1, and Pikm-1 HMA domain surface expression and interaction with Avr-Pik variants shown in bivariate contour plots. Clusters of cells that do not express the target protein (*11*) do not exhibit Avr-Pik binding. **c**, Flow cytometry quantification of Pikp-1 and Pikm-1 HMA domain interaction with Avr-Pik variants shown in univariate histograms. **d**, Exemplary flow cytometry ancestry plots showing the gating strategy applied to histograms in this study, as described in the Methods section. Plots from Pikm-1 HMA domain interaction with 1μM Avr-PikF are shown as an example.

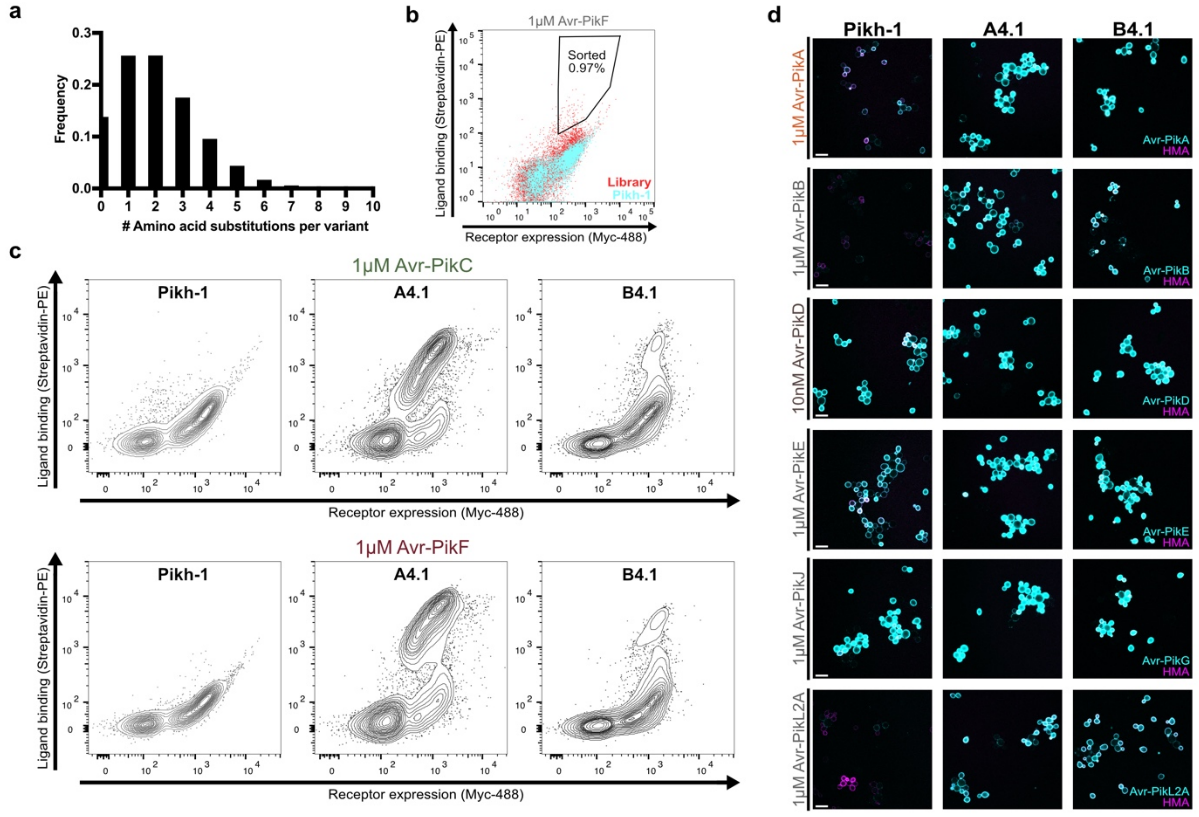


**e**

```
  Avr-PikA: MRVTTFNTFLLTLGTVAVVNAETGNKYIEKRAIDLSRERDPNFFDNADIPVPECFWFMFKNNVRQDAGTCYSSWKMDMKVGPNWVHIKSDDNCNLSGDFPPGWIVLGKKRPGF
  Avr-PikB: MRVTTFNTFLLTLGTVAVVNAETGNKYIEKRAIDLSRERDPNFFDNADIPVPECFWFMFKNNVRQDAGTCYSSWKMDIKVGPNWVHIKSDDNCNLSGDFPPGWIVLGKKRPGF
  Avr-PikC: MRVTTFNTFLLTLGTVAVVNAETGNKYIEKRAIDLSRERDPNFFDNPGIPVPECFWFMFKNNVRQDDGTCYSSWKMDMKVGPNWVHIKSDDNCNLSGDFPPGWIVLGKKRPGF
  Avr-PikD: MRVTTFNTFLLTLGTVAVVNAETGNKYIEKRAIDLSRERDPNFFDHPGIPVPECFWFMFKNNVRQDAGTCYSSWKMDMKVGPNWVHIKSDDNCNLSGDFPPGWIVLGKKRPGF
  Avr-PikE: MRVTTFNTFLLTLGTVAVVNAETGNKYIEKRAIDLSRERDPNFFDNPGIPVPECFWFMFKNNVRQDAGTCYSSWKMDMKVGPNWVHIKSDDNCNLSGDFPPGWIVLGKKRPGF
  Avr-PikF: MRVTTFNTFLLTLGTVAVVNAETGNKYIEKRAIDLSRERDPNFFDNADIPVPECFWFMFKNNVRQDAGTCYSSWKMDKKVGPNWVHIKSDDNCNLSGDFPPGWIVLGKKRPGF
  Avr-PikJ: MRVTTFNTFLLTLGTVAVVNAETGNKYIEKRAIDLSRERDPNFFDHPDIPVPECFWFMFKNNVRQDAGTCYSSWKMDMKVGPNWVHIKSDDNCNLSGDFPPGWIVLGKKRPGF
Avr-PikL2A: MRLVTFNTVFLTLGTISIVNAQNEYLDAKKHGIDLSRERAPNFVDHPGIPPSDCFWFLYKNYVRQDAGVCQSDWSFDMKIGQYWVTIHTDEGCRLSGIIPAGWLILGIKRLGF
            **:.****.:*****:::***:.     :*:.******* ***.*: .**  :****::** **** *.* *.*.:* *:*  ** *::*:.*.*** :* **::** ** **

  AvrPiz-t: MQFSTIITVCLFTGLASASFVQCNHHLLYNGRHWGTIRKKAGWAVRFYEEKPGQPKRLVAICKNASPVHCNYLKCTNLAAGFSAGTSTDVLSSGTVGSIGNDPQAQRQ
```

**Extended Data Fig. 2. Iterative selection leads to Pik-1 HMA domain variants with improved affinity for Avr-PikC and Avr-PikF. a**, Frequency of variants harboring the specified number of substitutions over the 78-amino acid HMA domain in the library of $2.2\times10^7$ Pikh-1 HMA domain variants. **b**, Bivariate dot plot of Pikh-1 HMA domain surface expression and interaction with 1µM Avr-PikF showing an exemplary FACS selection window. Cyan dots indicate cells expressing the wildtype Pikh-1 HMA domain, and red dots indicate cells expressing Pikh-1 HMA domain variants in the library. **c**, Flow cytometry quantification of Pikh-1, A4.1, and B4.1 HMA domain surface expression and interaction with 1µM Avr-PikC or Avr-PikF shown in bivariate contour plots. **d,** Immunofluorescence images of yeast cells expressing Pikh-1, A4.1, or B4.1 HMA domain in the presence of each effector at specified concentrations. Merged images show effector binding (cyan) and expression (magenta) of HMA domains. Scale bar, 10µm. **e**, Amino acid sequences of Avr-Pik and AvrPiz-t effectors from *Magnaporthe oryzae*.

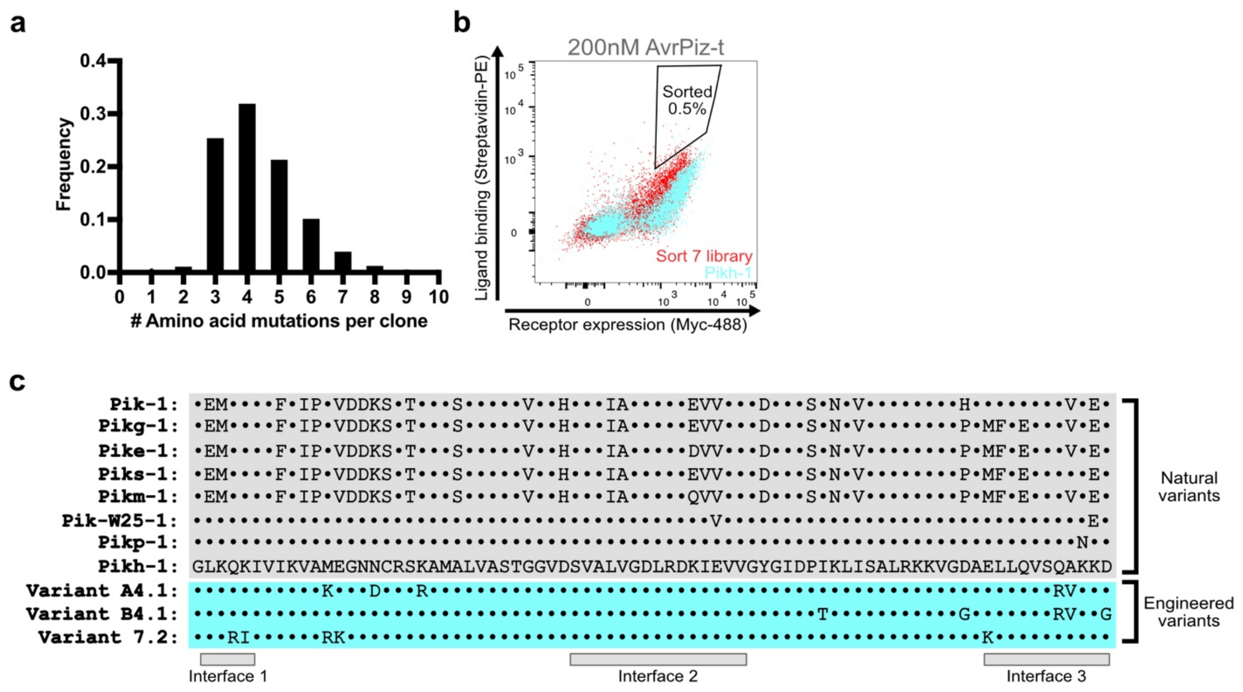


**Extended Data Fig. 3. Characterization of the second library and sequences of natural and engineered Pik-1 variants**. **a**, Frequency of variants harboring the specified number of substitutions over the 78-amino acid HMA domain in the library of 2.1×10$^{7}$ Pikh-1 HMA domain variants following the second round of mutagenesis. **b,** Bivariate dot plot of Pikh-1 HMA domain surface expression and interaction with 200nM AvrPiz-t showing an exemplary FACS selection window. Cyan dots indicate cells expressing the wildtype Pikh-1 HMA domain and red dots indicate cells expressing Pikh-1 HMA domain variants in the library following second round of mutagenesis. **c**, Amino acid sequences of Pik-1 HMA domains from rice alleles Pik-1, Pikg-1, Pike-1, Piks-1, Pikm-1, Pik-W25-1, Pikp-1, and Pikh-1 and engineered variants A4.1, B4.1, and 7.2. Interfaces known to interact with Avr-Pik are indicated below (*12–15*).

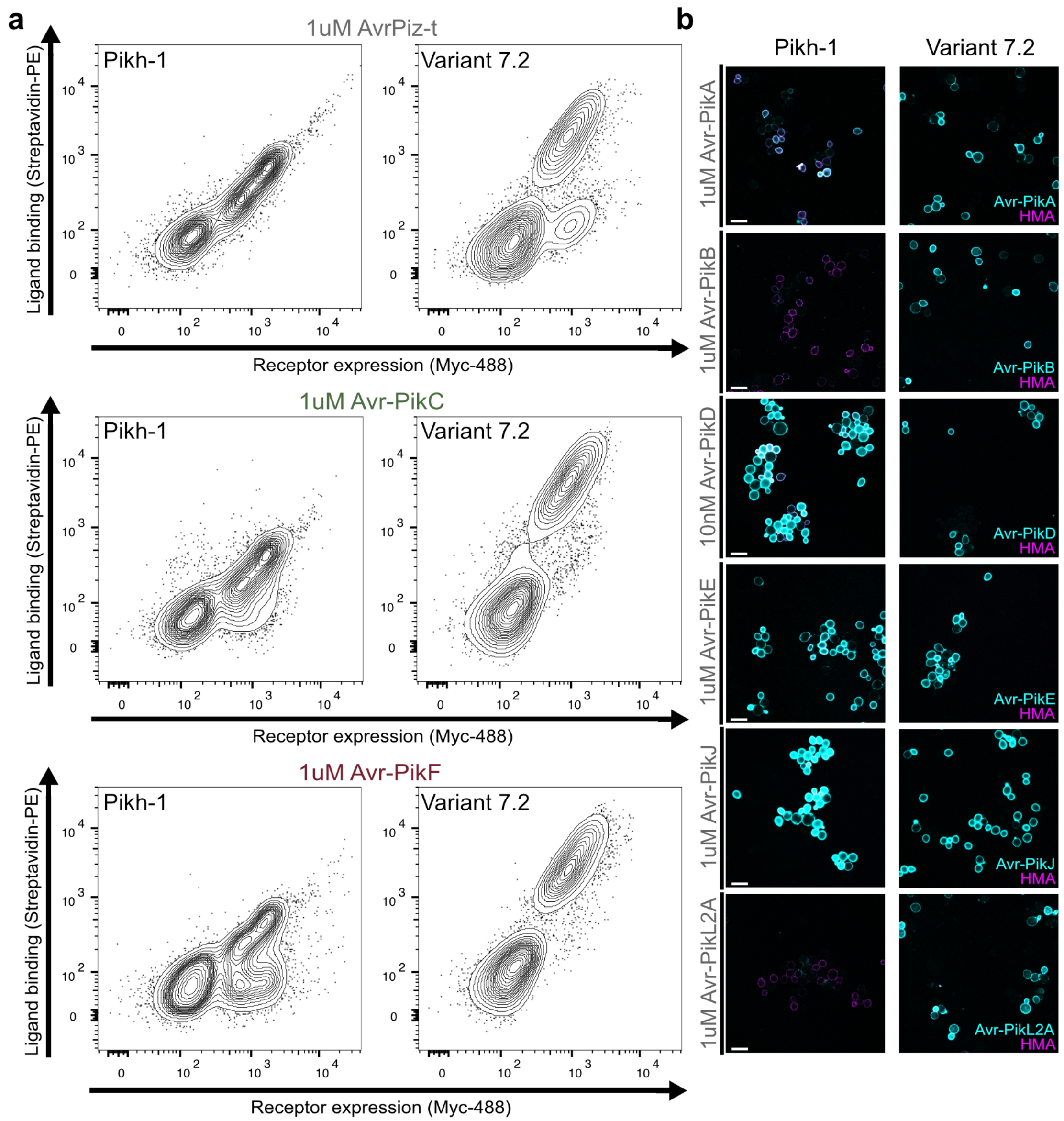


**Extended Data Fig. 4. Additional rounds of selection lead to a Pik-1 HMA domain variant with affinity for Avr-Pik and AvrPiz-t. a**, Flow cytometry quantification of Pikh-1 and 7.2 HMA domain surface expression and interaction with 1μM AvrPiz-t, Avr-PikC, or Avr-PikF shown in bivariate contour plots. **b**, Immunofluorescence images of yeast cells expressing Pikh-1 or 7.2 HMA domains in the presence of each effector at specified concentrations. Merged images show effector binding (cyan) and expression (magenta) of HMA domains. Scale bar, 10μm.

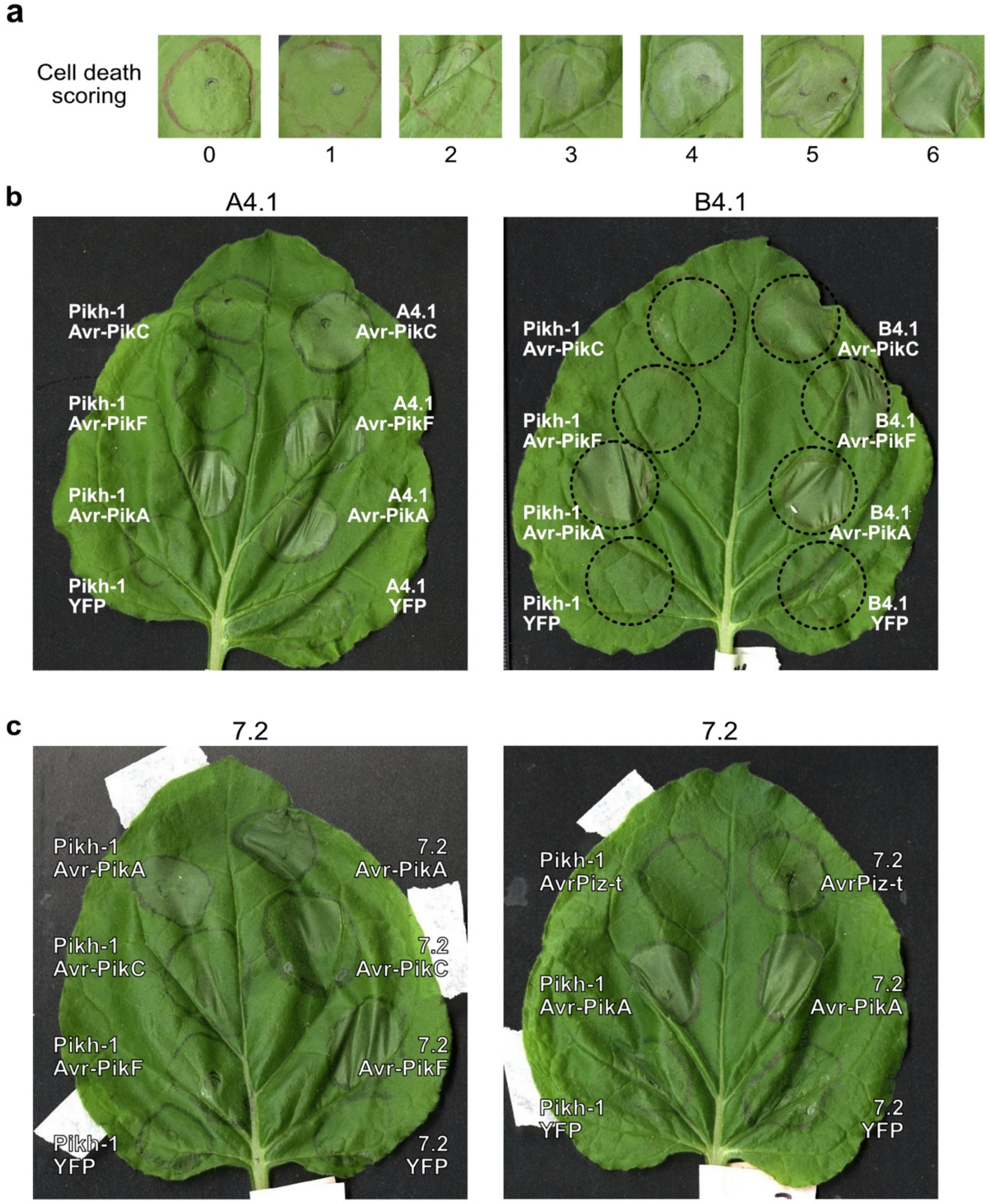


**Extended Data Fig. 5. Engineered Pik-1 immune receptors induce cell death response in *N. benthamiana*.** **a**, Representative images of *N. benthamiana* cell death scores on the 0-6 scale used in this study. **b**, Representative images of cell death assay. Pikh-1 harboring wildtype, A4.1, or B4.1 HMA domain was co-expressed with Avr-PikC, Avr-PikF, Avr-PikA (positive control), or YFP (negative control). **c**, Representative images of cell death assay. Pikh-1 harboring wildtype or 7.2 HMA domain was co-expressed with Avr-PikC, Avr-PikF, AvrPiz-t, Avr-PikA (positive control), or YFP (negative control).
